# An atypical ABC transporter is involved in antifungal resistance and host interactions in the pathogenic fungus *Cryptococcus neoformans*

**DOI:** 10.1101/2022.03.28.486166

**Authors:** Christopher J. Winski, Yuanyuan Qian, Shahriar Mobashery, Felipe H. Santiago-Tirado

## Abstract

ATP-binding cassette (ABC) transporters represent one of the largest protein superfamilies. Functionally diverse, ABC transporters have been implicated in many aspects of microbial physiology. The genome of the human fungal pathogen *Cryptococcus neoformans* encodes 54 putative ABC transporters and the majority of them remain uncharacterized. In a previous genetic screen for fungal regulators of phagocytosis, we identified an uncharacterized gene, CNAG_06909, that modulates host interactions. This gene encodes a half-size ABC transporter of the PDR-type, and phenotypic studies of a strain with this gene deleted revealed an altered antifungal susceptibility profile, including hypersensitivity to fluconazole (FLC). This gene, which we have named *PDR6*, localizes to the endoplasmic reticulum (ER) and plasma membrane (PM), and when absent, less ergosterol is observed in the PM. Additionally, we observed that the *pdr6*Δ strain displays a reduction in secreted polysaccharide capsular material. These changes to the cellular surface may explain the observed increased uptake by macrophages and the reduced intracellular survival. Finally, studies in mice demonstrate that Pdr6 function is required for normal progression of cryptococcal infection. Taken together, this study demonstrates a novel dual role for PDR transporters in *C. neoformans*, which could represent a potential target for antifungal therapeutics. Furthermore, the atypical half-size transporter encoded by *PDR6* is conserved in many fungal pathogens, but absent in model non-pathogenic fungi. Hence, this study provides for the first time, a function for this unique group of fungal half-size PDR transporters that, although conserved, remain largely understudied.

**IMPORTANCE:** Conserved across all kingdoms of life, ABC transporters comprise one of the largest protein families. They are associated with multidrug resistance, affecting aspects such as resistance to antimicrobials or anti-cancer drugs. Despite their importance, they are understudied in fungal pathogens. In the environmental fungus *Cryptococcus neoformans*, a leading cause of fungal infections, only a few ABC transporters have been studied. Here we characterize an atypical, half-size, ABC transporter of the PDR-type, that affects both antifungal resistance and host-pathogen interactions. PDR-type transporters are only present in fungi and plants, and this subgroup of half-size transporters is conserved in fungal pathogens, yet their function was completely unknown. Because the current treatments for cryptococcal infection are suboptimal, understanding the mechanisms of antifungal resistance and the host interactions that drive the infection is critical to improve the management of this disease. Here we provide insights into these important aspects of cryptococcal pathogenesis.

## INTRODUCTION

Present in every living organism, ATP-binding cassette (ABC) transporters constitute a large superfamily of integral membrane proteins. Functionally, ABC transporters are responsible for the transport of diverse substrates across membranes, but have also been implicated in mRNA translation and peroxisome biogenesis (1, 2). Structurally, ABC transporters contain at least one nucleotide-binding domain (NBD) and one transmembrane domain (TMD), both of which are evolutionary conserved. Most ABC transporters have a domain arrangement of TMD1-NBD1-TMD2-NBD2 ([TMD-NBD]_2_) (3). The orientation and molecular architecture of these domains is used to organize ABC transporters into subgroups. The pleiotropic drug resistance (PDR) subgroup, which is present in only fungi and plants, has a reverse configuration [NBD-TBD]_2_ (4, 5). Identified and predominantly studied in the non-pathogenic yeast *Saccharomyces cerevisiae*, they remain understudied in pathogenic fungi. This is despite the abundance of fungal PDR proteins and the fact that fungal infections globally kill close to 1.6 million people yearly (6, 7). The few that have been studied mostly act as efflux pumps for xenobiotics, including antifungal drugs. Nonetheless, the sheer number and diversity of PDR genes found in fungal genomes suggests a broad involvement in cellular processes. Moreover, there is a subgroup of half-size PDR transporters (containing only one NBD and TMD) whose function remains unknown. Because all functional PDR transporters studied so far have the typical [NBD-TBD]_2_ domain arrangement, these half-size ABC transporters with a [NBD–TMD] topology have been left out of PDR protein phylogenetic studies (4, 5, 8).

*Cryptococcus neoformans* is an environmental, opportunistic fungal pathogen responsible for over 180,000 deaths annually in the HIV+ population alone (9, 10). Upon inhalation and subsequent lung colonization, *C. neoformans* can disseminate to the central nervous system (CNS) resulting in cryptococcal meningitis, with a mortality rate as high as 81% (10). The high mortality rate is a direct consequence of the complexities associated with cryptococcal pathogenesis coupled with a paucity of effective antifungals. Currently, the most effective treatment for cryptococcal meningitis includes a combinatorial regimen of amphotericin B (AMB) and 5-flucytosine (5-FC). AMB is a fungicidal polyene that targets ergosterol, an essential lipid in the fungal plasma membrane (PM). The pyrimidine 5-FC disrupts DNA/RNA synthesis (11). Notwithstanding toxicity associated with these agents, they are also costly, which makes them less available to resource-poor countries, where the need is highest. Instead, patients in these regions are treated with the less effective fluconazole (FLC) monotherapy. FLC is a fungistatic azole that inhibits lanosterol 14α-demethylase, an enzyme in the ergosterol biosynthesis pathway encoded by *ERG11* (12). FLC monotherapy has not only been implicated in higher relapse rates, but several recent studies have reported an increase in the incidence of FLC resistance in *C. neoformans*, making the treatment of cryptococcal meningitis even more challenging (13–16).

Several FLC resistance mechanisms have been reported in *C. neoformans*. These vary from mutations in the *ERG11* gene to heteroresistance due to genome plasticity including polyploidy, aneuploidy, loss of heterozygosity, and copy-number variation (17–20). Additionally, the overexpression of ABC efflux pumps has been implicated in FLC resistance (21). There are 54 putative ABC transporters in the *C. neoformans* genome and only a few have been characterized. The roles of *C. neoformans AFR1*, *PDR5* (also called *AFR2*), and *MDR1* as azole efflux pumps have been well documented, where *AFR1* functions as the major pump and *AFR2* and *MDR1* provide additive roles in the management of FLC stress (22–25). It is evident that the function of most of the ABC transporters encoded in the *C. neoformans* genome remain unknown. In an earlier report on fungal regulators of phagocytosis, we identified an uncharacterized gene, CNAG_06909, as a modulator of the fungal-host interaction (26). In the present study, we demonstrate that CNAG_06909 encodes a half-size PDR-transporter and that disruption of the gene results in hypersensitivity to FLC. Following nomenclature guidelines for the field (27), we have designated the gene as pleiotropic drug resistance-6 (*PDR6*) because it does not have a direct ortholog in the model yeast *S. cerevisiae* and the molecular architecture is consistent with the PDR subgroup of ABC transporters (see Supplemental Text S1 for more details). We show that Pdr6 does not function as a FLC efflux pump but its deletion results in decreased levels of ergosterol in the PM, accompanied by mild thermosensitivity and decreased secretion of capsular material. These phenotypes are unique to *PDR6* as they were not observed in the previously characterized ABC transporters. Furthermore, the *pdr6*Δ strain exhibits diminished intracellular survival *in vitro* and is attenuated in a murine model of infection. When the gene is expressed in model yeast, it localizes to the ER and PM, consistent with a role in ergosterol synthesis/transport. Notably, this study provides for the first time a function for the group of unconventional, yet conserved, half-size PDR transporters, which has remained largely obscure. These studies highlight the importance of fungal PDR-type transporters and broaden our understanding of the specialization of ABC transporters in microbial pathogens.

## RESULTS

### *C. neoformans PDR6* is a member of a conserved group of half-size PDR transporters

Present in only fungi and plants, PDR-type ABC transporters represent a unique subset of proteins primarily implicated in drug resistance and detoxification (4). Although fungal PDR-type transporters closely resemble mammalian ABCG transporters, which are half-size, only the full-size fungal PDR proteins have been analyzed and studied (5, 8, 28). Given that *C. neoformans PDR6* is half-size, we performed a phylogenetic analysis to determine how widespread *PDR6* sequences are in the fungal kingdom (Fig. 1A). All fungi belonging to the Pezizomycotina subphylum and all the pathogenic fungi analyzed have a *PDR6* ortholog, implicating *PDR6* in fungal pathogenesis in many subphyla. Regardless of the presence of a *PDR6* ortholog, all the PDR-type ABC transporter genes from the organisms in Figure 1A were used to construct a global phylogenetic tree (Fig. 1B). Analysis of the fungal PDR genes showed the presence of seven major clades. Clade I is the largest and includes *S. cerevisiae PDR5*, *PDR10*, *PDR15*, and *C. neoformans PDR5* (hereafter referred by its alias *AFR2)*. Clade II is the smallest and contains sequences that are homologous to *S. cerevisiae PDR12*, *PDR18* and *SNQ2.* Clade III sequences are absent in Saccharomycotina, but contain *C. neoformans AFR1*. *S. cerevisiae PDR11* and *AUS1* represent the genes in clade IV. Clades I, II, III, IV, and VIa all contain full-length *PDR* genes, whereas clades V and VIb represent atypical groups of PDR genes that only encode one nucleotide-binding domain (NBD) and one transmembrane domain (TMD). *S. cerevisiae ADP1* is representative of the genes in clade V that all contain a NBD and TMD with an additional unique sequence (most commonly an EGF domain). *C. neoformans PDR6* and the eleven other half-size PDR genes in clade VIb represent a completely uncharacterized group of fungal genes with no known function. Interestingly, the fungi represented in clade VIb are all plant or animal pathogens, again suggestive that *PDR6* and its orthologs may be implicated in fungal virulence. Topology prediction by CAMPS (29) shows that *C. neoformans PDR6* encodes a 626 amino acid half-size transporter, with a single NBD containing characteristic Walker A, Walker B, and ABC motif features followed by a single TMD, composed of six transmembrane helices (Fig. 1C). These analyses show that *C. neoformans PDR6* encodes a half-size PDR-type ABC transporter that is evolutionarily conserved, with orthologs present in all fungal pathogens that we analyzed, and contains all the features of a functional ABC transporter.

**FIG 1.**
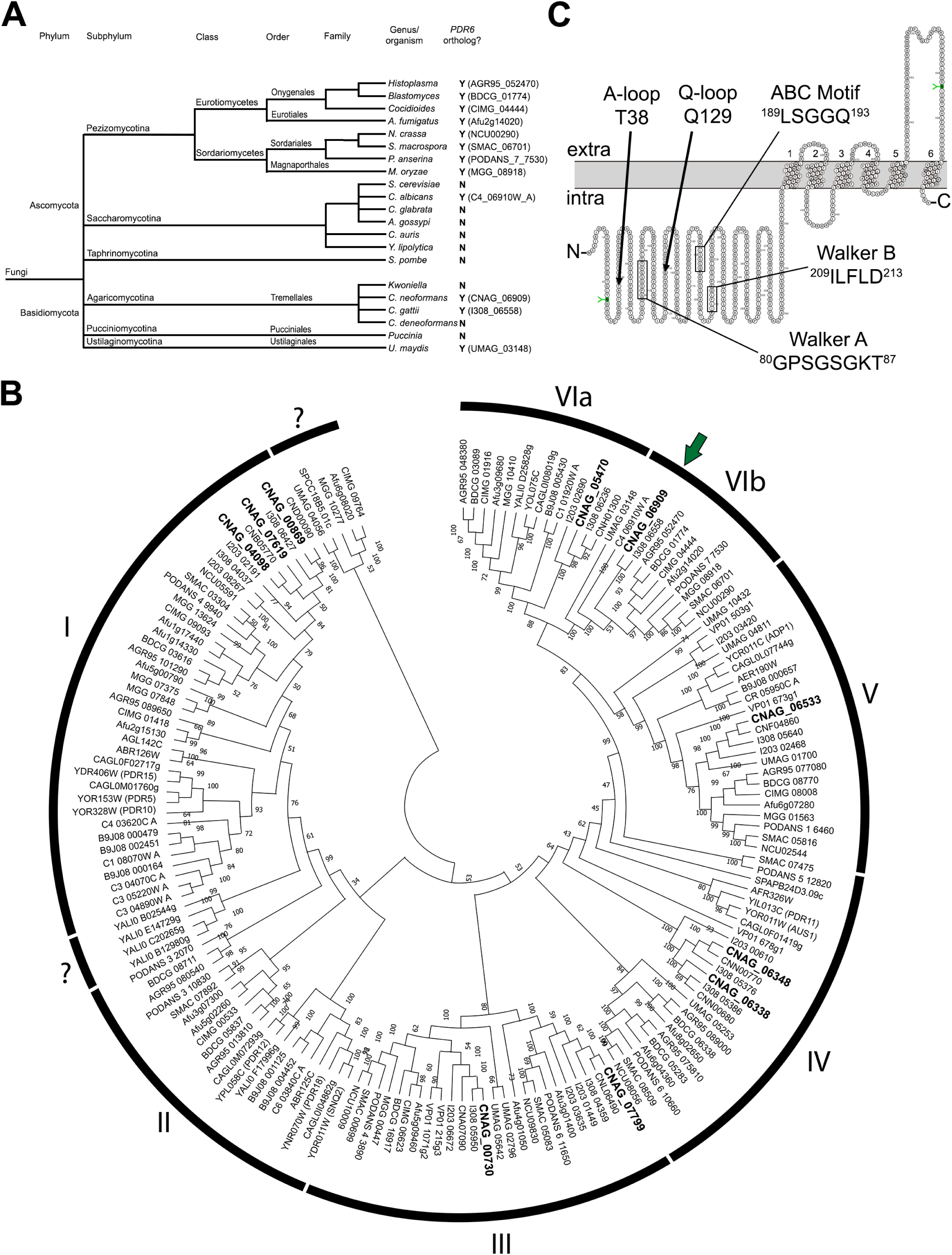
*PDR6* contains all the features of a PDR-type ABC transporter and is conserved in a subset of fungal pathogens. (A) Cladogram showing the fungal distribution of *PDR6* orthologs in the two main phyla. When present, the gene name of the ortholog is stated within parentheses. (B) Global phylogenetic tree showing the evolutionary relationships of PDR-type ABC transporters in 21 fungal organisms. Gene names are given in NCBI nomenclature. Tree was generated by Maximum Likelihood Method using MEGA-X and confidence values determined from 500 bootstraps with bootstrap scores shown at all nodes. The tree shown is the bootstrap consensus tree. Analysis reveals VI major clades with *C. neoformans PDR6* housed in clade VIb (denoted by dark green arrow). The 10 PDR-type transporters in *C. neoformans* are in bold font and are listed in Supplementary Text S1. (C) Topology prediction of Pdr6 protein by Computational Analysis of the Membrane Protein Space (CAMPS) software showing the domain arrangement of the half-size PDR transporter. The 5 main consensus motifs of the nucleotide binding domain (NBD) are depicted. The residues in green with a green ‘Y’ are predicted N-glycosylation sites.

### Deletion of *PDR6* affects resistance to azoles and the drug profile of other antifungals

Given that cryptococcal PDR-type ABC transporters Afr1 and Afr2 have been implicated in the efflux and resistance of antifungal compounds (23–25), we hypothesized that Pdr6 would function in a similar manner. To assess the function/specificity of Pdr6, we used a *C. neoformans* strain lacking the *PDR6* gene (*pdr6*Δ) and tested its growth on FLC gradient plates and Etest strips (Fig. 2A), and performed broth microdilution assays to measure the minimal-inhibitory concentrations (MICs) of several related compounds (Fig. 2B-G and Table 1). Compounds included five azole-class antifungals (FLC, itraconazole (ITC), voriconazole (VRC), ketoconazole (KTC), and posaconazole (PSC)), one polyene-class antifungal (amphotericin B (AMB)), one DNA/RNA synthesis inhibitor (5-fluorocytosine (5-FC)), one protein-synthesis inhibitor (cycloheximide (CHX)), one antineoplastic agent (nocodazole), two other xenobiotics (berberine (BER), and paraquat (PQ)), one superoxide generator (menadione), and one echinocandin-class antifungal (caspofungin (CSF)). In addition to the *pdr6*Δ mutant strain, we used the *afr1*Δ, *afr1*Δ/*afr2*Δ/*mdr1*Δ, and *erg3*Δ strains as controls, since all have displayed increased FLC susceptibility (24, 30).

**FIG 2.**
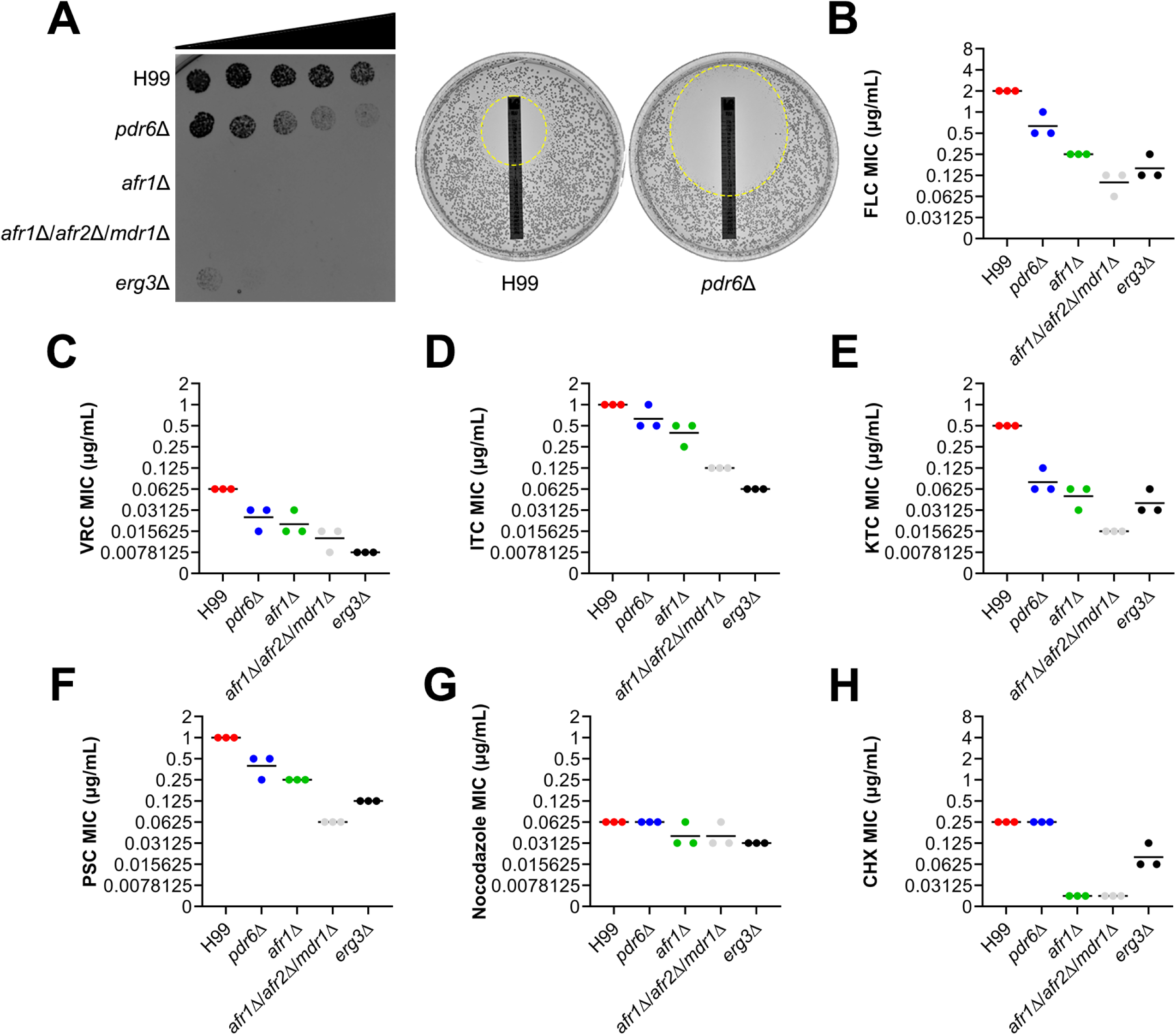
*PDR6* is involved in azole resistance. (A) The *pdr6*Δ mutant shows increased sensitivity to FLC gradient plates and Etest strips. (Left) 500 cells of the indicated strains were spotted across a RPMI-1640 agar plate with a gradient of FLC concentration (denoted by black triangle). The gradient goes from near zero at the left side to a peak concentration of 4 μg/mL at the right side of the plate. (Right) FLC Etest strips were applied to RPMI-1640 agar plates with 2 x 10^5^ cells of the indicated strain. The zone of inhibition is highlighted with a dashed circle, and the MICs of H99 and the *pdr6*Δ mutant were 6 and 0.38 μg/mL, respectively. These plates were incubated for 72 hr at 30°C. (B-H) Quantitative broth microdilution assays were used to compare MICs of several antifungal compounds in the WT and the indicated mutant strains. The CLSI M27-A3 reference method was repeated three times for each compound and the bars represent the average of the MICs. The details for each strain’s MICs are listed in Table 1. FLC, fluconazole; VRC, voriconazole; ITC, itraconazole; KTC, ketoconazole; PSC, posaconazole; CHX, cycloheximide.

**TABLE 1.** MICs of antifungals and xenobiotics.^a^

| Strain | MICs (µg/mL) |  |  |  |  |  |  |  |  |  |  |  |  |  |
| --- | --- | --- | --- | --- | --- | --- | --- | --- | --- | --- | --- | --- | --- | --- |
|  | FLC | ITC | VRC | KTC | PSC | AMB | 5-FC | CHX | NOC | BER | PQ | TSA | MEN | CSF |
| **H99** | 2 | 1 | .0625 | 0.5 | 1 | 4 | 2 | 0.25 | 0.0625 | 8 | 12.5 | 2 | 16 | 32 |
| ***pdr6Δ*** | 0.5 – 1 | 0.5 – 1 | 0.0156<br>–<br>0.0312 | 0.0625<br>–<br>0.125 | 0.25 –<br>0.5 | 4 | 4 | 0.25 | 0.0625 | 8 | 12.5 | 2 | 16 | 64 |
| ***afr1Δ*** | 0.25 | 0.25 –<br>0.5 | 0.0156<br>–<br>0.0312 | 0.0312<br>–<br>0.0625 | 0.25 | 4 | 2 | 0.0156 | 0.0312<br>–<br>0.0625 | 8 | 12.5 | 0.5 | 16 | 64 |
| ***afr1Δ/***<br>***afr2Δ/***<br>***mdr1Δ*** | 0.0625<br>–<br>0.125 | 0.125 | 0.0078<br>–<br>0.0156 | 0.0156 | 0.0625 | 4 | 4 | 0.0156 | 0.0312<br>–<br>0.0625 | 8 | 12.5 | 0.5 | 8 | 64 |
| ***erg3Δ*** | 0.125<br>– 0.25 | 0.0625 | 0.0078 | 0.0312<br>–<br>0.0625 | 0.125 | 4 | 2 | 0.0625<br>–<br>0.125 | 0.0312 | 0.5 | 3.125 | 0.5 | 16 | 32 |

<sup>a</sup> b Values given are the range of MICs obtained from at least three independent experiments. FLC, fluconazole; ITC, itraconazole; VRC, voriconazole; KTC, ketoconazole; PSC, posaconazole; AMB, amphotericin B; 5-FC, 5-fluorocytosine; CHX, cycloheximide; NOC, nocodazole; BER, berberine; PQ, paraquat; TSA, trichostatin A; MEN; menadione; CSF, caspofungin.

Among the compounds tested, we found that the *pdr6*Δ mutant displayed a hypersensitive phenotype specifically towards the azole class antifungals, albeit not as drastic as the *afr1*Δ, *afr1*Δ/*afr2*Δ/*mdr1*Δ, or *erg3*Δ strains (Fig. 2 and Table 1). The MICs of the non-azole compounds in the *pdr6*Δ strain were comparable to those of the H99 WT, suggesting a specific role of Pdr6 in azole resistance. Additionally, previous studies have shown a relationship between environmental pH and antifungal efficacy (31), so we performed FLC susceptibility assays at alkaline and acidic conditions (Fig. S1A,B). Regardless of the pH, the MICs of the WT and *pdr6*Δ strains displayed similar trends indicating that *PDR6* functions in a pH-independent manner.

Given the increase in azole susceptibility by the *pdr6*Δ mutant, we performed checkerboard assays with four common antifungals, FLC, AMB, 5-FC, and CSF (Fig. 3). Interestingly, we found that FLC coupled with CSF functions differently in the *pdr6*Δ strain when compared to the WT. The combination works strongly in an additive manner in the *pdr6*Δ strain but shows minimal combinatorial effects in the WT. This finding was unexpected given that CSF is ineffective at treating cryptococcal infections. Additional experimentation is required to explain the observed finding. Nonetheless, these data show that *PDR6* is involved in azole resistance, is not affected by pH, and can alter antifungal combinatorial behaviors. All of these results were confirmed using three independently recreated *pdr6*Δ mutants (Fig. S1C and Materials and Methods).

**FIG 3.**
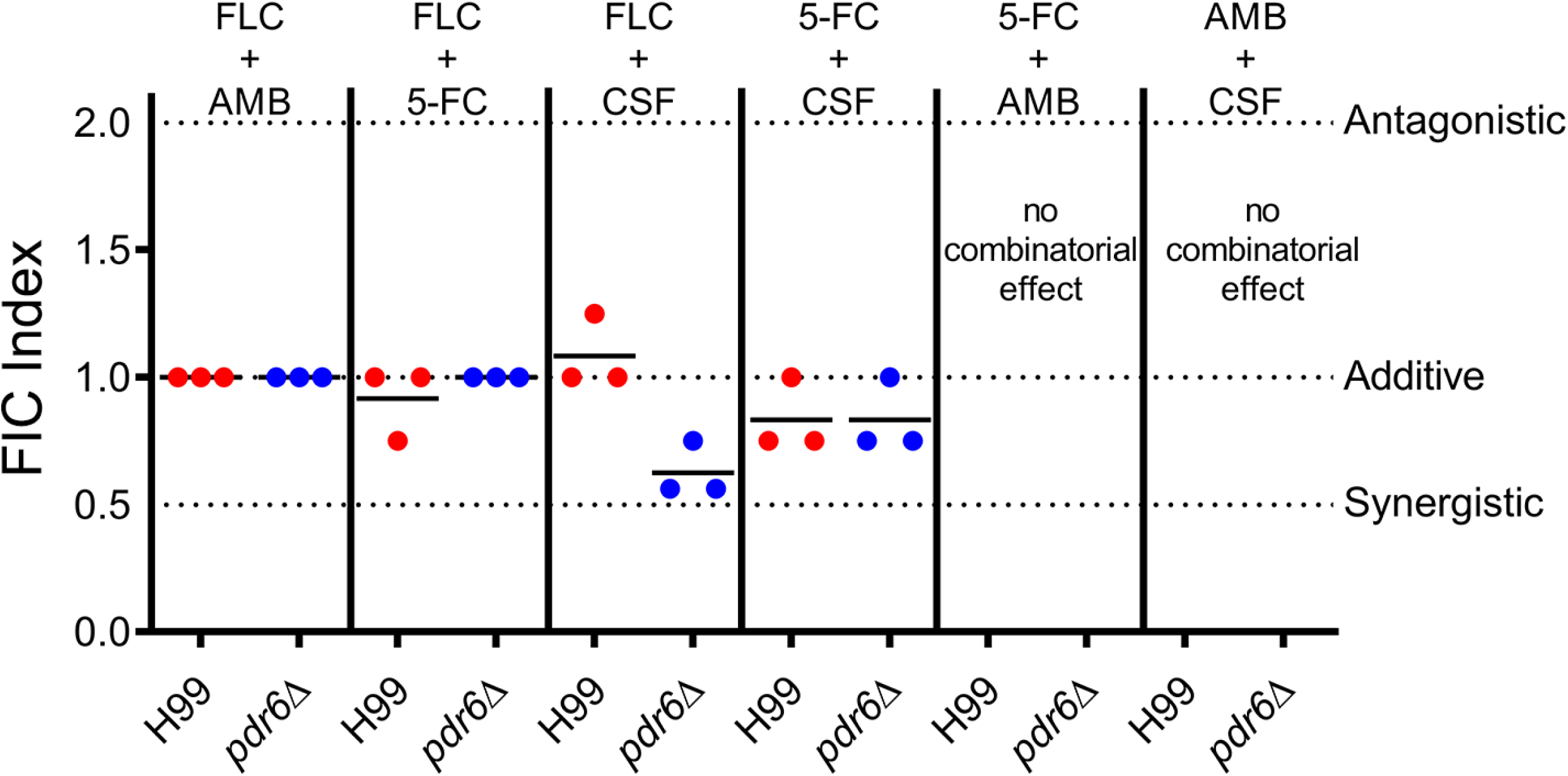
Combinatorial FLC + CSF treatment have an additive effect in *pdr6*Δ mutant. Checkerboard assays assessed changes in drug combinatorial activity in H99 and *pdr6*Δ strains. The combination of FLC + CSF resulted in average sums of FIC of 1.083 and 0.625 in the H99 and *pdr6*Δ strains, respectively. Calculations indicate an additive effect trending into synergistic in the absence of *PDR6*. Checkerboard assays were repeated three times for each combination and black bars represent the average sum of FIC.

### Deletion of *PDR6* does not affect drug accumulation

The increased azole susceptibility of the *pdr6*Δ mutant led us to hypothesize that Pdr6 could function as an efflux pump, similar to many of the characterized fungal PDR transporters. Rhodamine 6G (R6G) and Nile Red (NR) are fluorescent xenobiotics that have been used as generic probes to measure ABC transporters’ efflux activity (32, 33). To determine if Pdr6 functions as an efflux pump, we measured the intracellular accumulation of R6G and NR by flow cytometry (Fig. 4A). Surprisingly, no difference was observed in the mean fluorescent intensity (MFI) values of either R6G or NR in the *pdr6*Δ mutant, compared to the WT. In contrast, the *afr1*Δ and *afr1*Δ/*afr2*Δ/*mdr1*Δ strains accumulated high levels of these xenobiotics. We reasoned that Pdr6 may not recognize R6G or NR and potentially, Pdr6 is strictly an azole-specific transporter. To test this, we synthesized a dansyl amide labeled-FLC molecule, termed Probe 1 (Fig. 4B, left), that has been used to visualize the intracellular localization of FLC in *Candida albicans* (34). Although the modification of FLC with dansyl amide affected its antifungal activity, the strains more sensitive to FLC were also more sensitive to Probe 1, indicating a similar mechanism of action (Fig. 4B, right). Consistently, the intracellular accumulation of Probe 1 was significantly increased in the *afr1*Δ and *afr1*Δ/*afr2*Δ/*mdr1*Δ strains (Fig. 4C). However, there was no difference in the intracellular accumulation of Probe 1 between the *pdr6*Δ strain and WT, suggesting that Pdr6 is not a major efflux pump for FLC and instead affects azole activity by a different mechanism. We also used light microscopy to assess the intracellular localization and accumulation/fluorescent intensity of Probe 1 between strains and no differences in location were observed, while fluorescent intensity values supported the findings above (data not shown). To provide definitive support that Pdr6 is not acting as a FLC-specific efflux pump, we directly measured accumulation of FLC using a liquid chromatography-mass spectrometry (LC-MS)-based FLC accumulation assay (Fig. 4D). We found that the *pdr6*Δ mutant once again displayed levels comparable to the WT, while the *afr1*Δ/*afr2*Δ/*mdr1*Δ positive control strain accumulated significant amounts of FLC. Taken together, these data support that Pdr6 is not functioning as a drug-efflux pump and that the hypersensitivity to azole antifungals in the *pdr6*Δ strain is not due to increased accumulation of these drugs.

**FIG 4.**
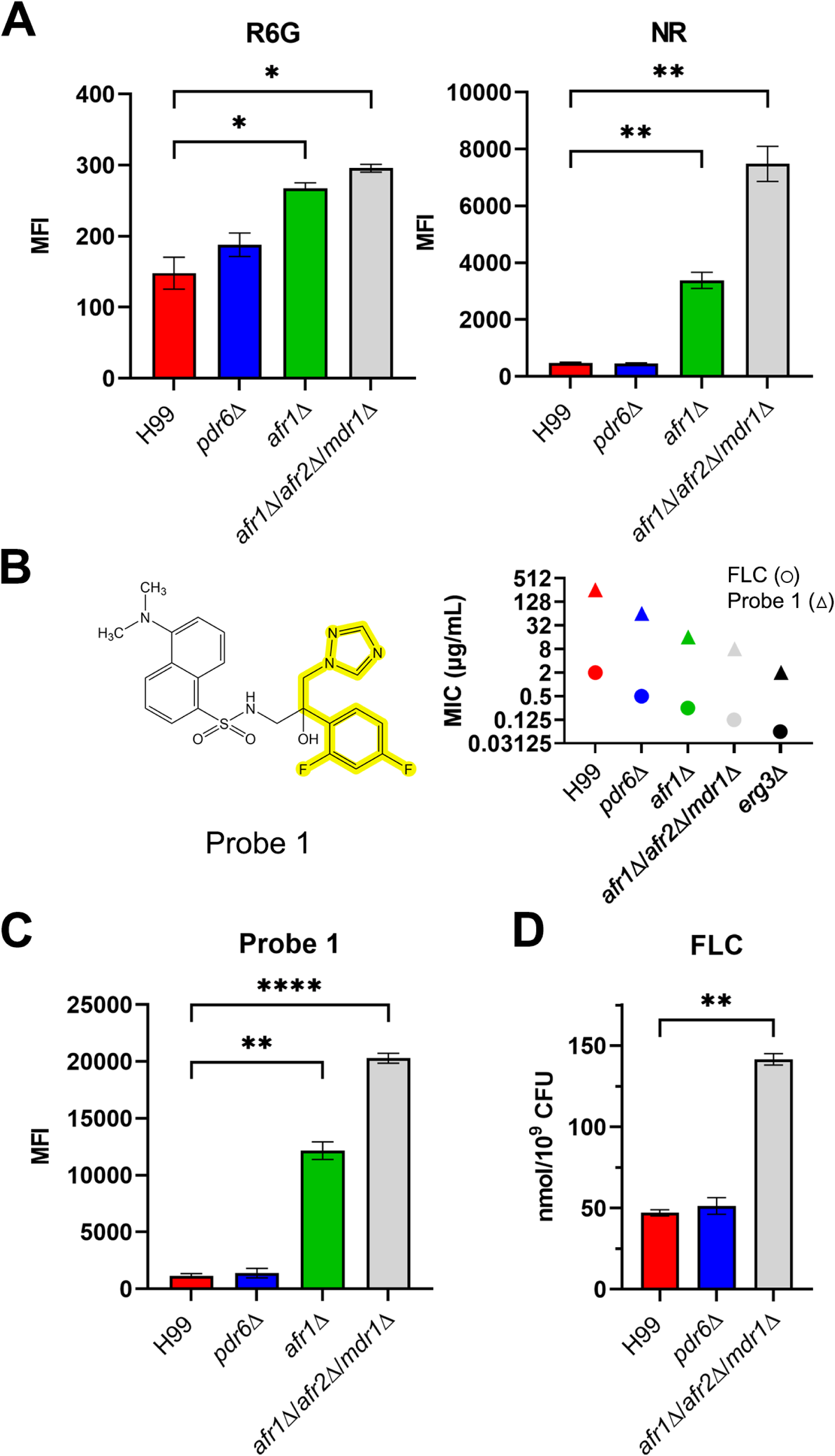
Deletion of *PDR6* does not affect intracellular accumulation of fluorescent reporters or of FLC. (A) H99 and indicated deletion strains were incubated with 10 μM R6G or 7 μM NR for 30 min at 30°C and accumulation of the reporter in the cells was analyzed by flow cytometry as described in Materials and Methods. (B) (Left) Chemical structure of Probe 1 with FLC backbone highlighted. (Right) Broth microdilution assays were performed to compare the MICs of FLC with Probe 1. Regardless of strain, a 128-fold increase was observed in the MICs of Probe 1 when compared to FLC. Microdilution assays were repeated three times and shown are the results from one representative experiment. (C) Indicated strains were incubated with 10 μg/mL of Probe 1 and accumulation assessed by flow cytometry as above. (D) Accumulation of FLC was assessed in H99 and deletion strains using LC-MS/MS. Strains were incubated with 16 μg/mL of FLC for 30 min at 30°C. All values represent the mean ± SEM from three to four independent experiments. Significance was determined using one-way ANOVA compared to H99 (Brown-Forsythe and Welch’s corrected); *P<0.05, **P<0.01, ****P<0.0001.

### *PDR6* is involved in ergosterol trafficking

To mechanistically explain Pdr6’s contribution to antifungal resistance, we next considered azoles’ mode of action. It is well documented that the target of azoles is Erg11, an enzyme in the ergosterol biosynthesis pathway (18, 35). Disruptions to the ergosterol metabolic pathway can result in accumulation of sterols, which have been implicated in both diminished and increased effectiveness of azole compounds (30, 36). We, therefore, hypothesized that the azole resistance phenotype might be a result of *Pdr6*’s involvement in ergosterol transport. To quantify ergosterol content in the cryptococcal PM, we used filipin, a sterol-binding fluorescent dye (Fig. 5). We found a significant difference in fluorescence between the WT and the *pdr6*Δ mutant. Specifically, the deletion of *PDR6* resulted in decreased fluorescence, suggesting that less sterol is present and that Pdr6 plays a role in either the synthesis of sterols or their trafficking; ABC transporters have been implicated in the latter (1, 37, 38), supporting a potential role for Pdr6 in ergosterol transport to the PM. Notably, neither the *afr1*Δ nor *afr1*Δ/*afr2*Δ/*mdr1*Δ strains exhibited reductions in filipin staining, indicating that the reduced sterol is specific for *pdr6*Δ. As a control, we used *erg3*Δ strain, which is expected to have low levels of ergosterol and, consistently, had almost no detectable staining. We further corroborated the quantitative microscopy results using a fluorescent plate reader, confirming that the *pdr6*Δ mutant has reduced overall ergosterol, compared to the WT (data not shown). These findings provide context for the previously observed antifungal phenotype and demonstrate that the *pdr6*Δ strain has an altered PM with less ergosterol, consistent with a role of *PDR6* in ergosterol transport/trafficking. Lastly, an indirect role of *PDR6* in azole resistance is supported by the fact that the presence of FLC does not affect the expression of *PDR6* (Fig. S2A), while growth under host-like conditions (DMEM, 37 °C, 5% CO_2_) induce a rapid and sustained increase in transcript levels (Fig. S2B). Ergosterol is essential for thermotolerance and survival inside mammalian hosts (38), so if *PDR6* is involved in ergosterol transport, then it would be expected to be overexpressed under host-like conditions.

**FIG 5.**
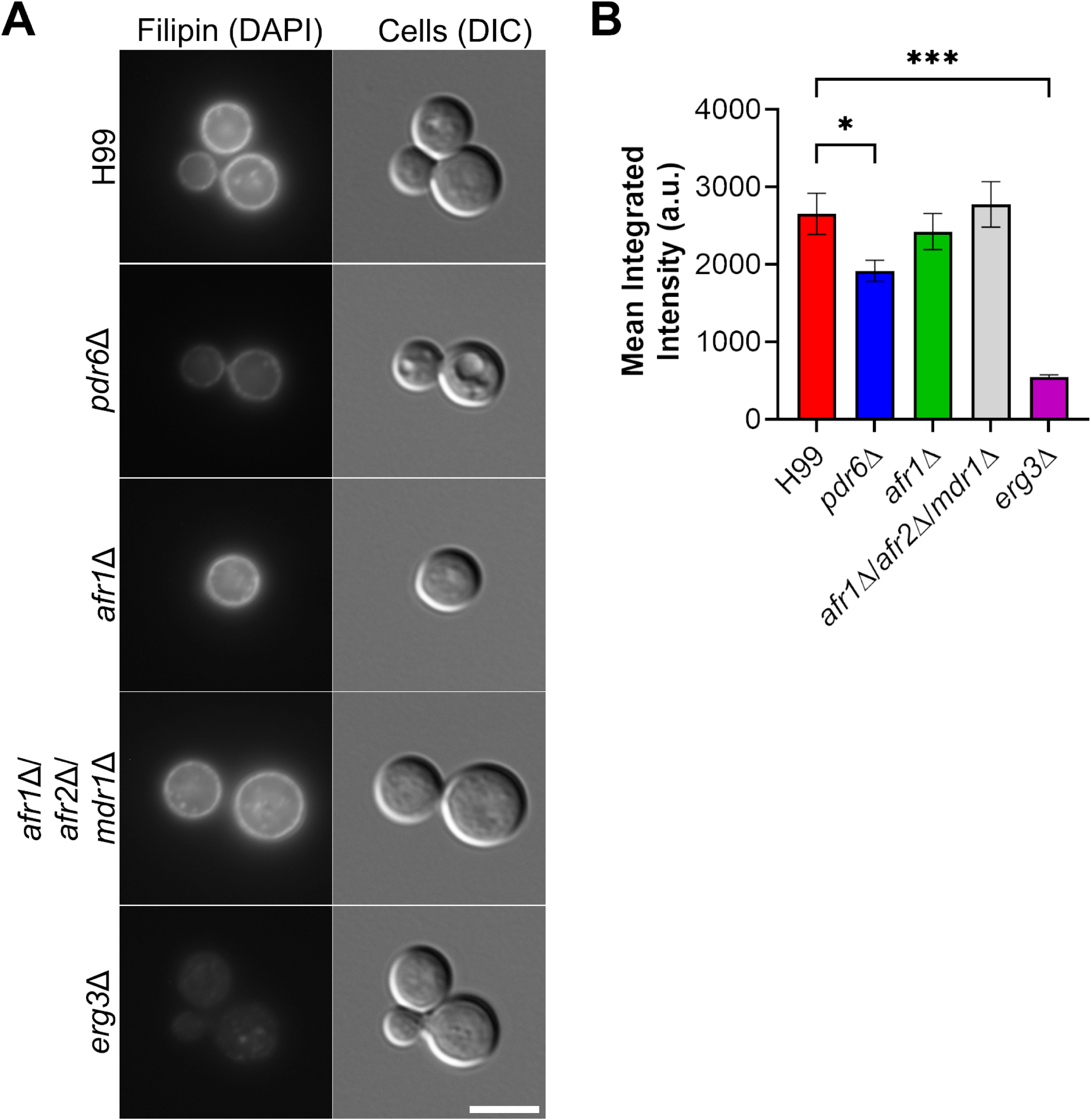
*pdr6*Δ mutant displays reduced sterol content. (Left) To assess sterol content in WT and other mutant strains, the cells were incubated with 5 μg/mL Filipin III for 5 min at 30°C and visualized microscopically. Scale bar, 5 μm. (Right) Quantification of fluorescence using a Cell Profiler pipeline. Values represented are the mean ± SEM of 3 to 4 independent experiments where n=50 cells per experiment. Significance was determined using one-way ANOVA with multiple comparisons (Brown-Forsythe and Welch’s corrected); *P<0.05, ***P<0.001.

### Pdr6 localizes to the ER and PM

To better understand the function of Pdr6, we generated a C-terminally tagged *PDR6*-GFP integrative construct under the control of the *PDR5* promoter and introduced it into the *S. cerevisiae* strain ADΔΔ (39, 40). This strain is missing seven major endogenous ABC transporters and expresses a mutant Pdr1 transcription factor, driving overexpression of PDR genes. This strain not only allows for the visualization of Pdr6 localization, but also a more direct assessment of efflux activity, provided cryptococcal Pdr6 would be the only major PDR protein in the cell. Visualization of the ADΔΔ+Pdr6-GFP strain revealed clear perinuclear and peripheral staining, characteristic of the yeast endoplasmic reticulum (ER) (Fig. 6A). Similarly, a control ADΔΔ strain expressing Afr1-GFP, a full-length PDR-type transporter, displayed comparable localization patterns consistent with the ER and PM (Fig. 6A). No GFP fluorescence was seen in the strain transformed with an empty vector (Fig. 6A).

**FIG 6.**
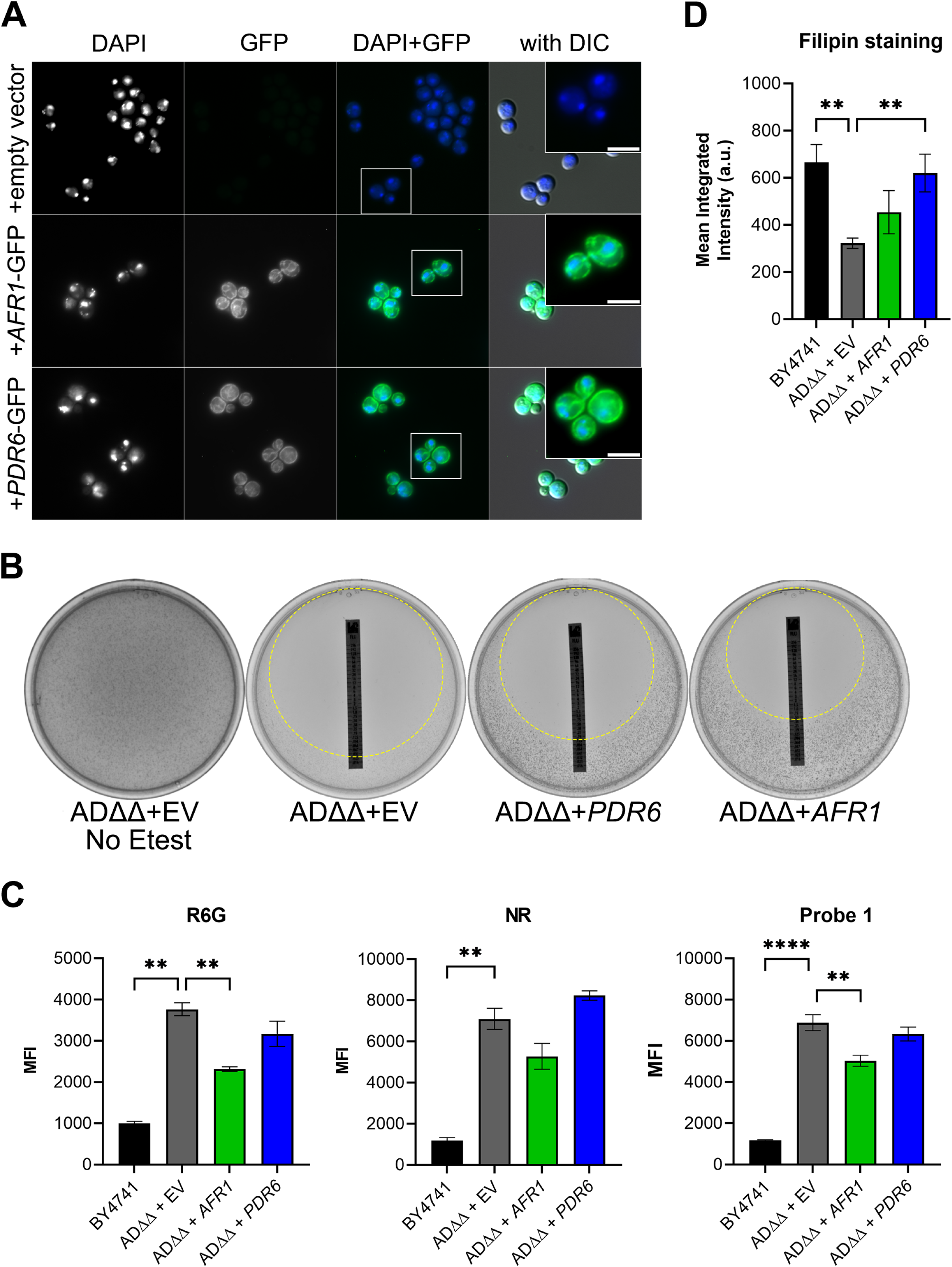
Pdr6 localizes to the ER and PM in *S. cerevisiae* but do not affect intracellular drug accumulation. (A) Visualization of *S. cerevisiae* ADΔΔ expressing *C. neoformans PDR6*-GFP, *AFR1*-GFP, or empty vector. DAPI staining was included to highlight the perinuclear staining. The GFP-tagged proteins localization is consistent with the ER and PM. Scale bar, 3.8 μm. (B) Growth of the various ADΔΔ strains was assessed using FLC Etest strips in RPMI plates. Strains expressing *PDR6* (MIC = 0.25 µg/mL) and *AFR1* (MIC = 0.75 µg/mL) display increased resistance to FLC compared to empty vector control (MIC = 0.064 µg/mL) but the strain expressing *AFR1* was most resistant. Plates were incubated at 30°C for 96 hr and imaged. The zones of inhibition are highlighted with a dashed circle. (C) Accumulation of the indicated reporters was assessed by flow cytometry as described in Fig. 4. There was no difference between the accumulated drugs in the empty vector control and the strain expressing *PDR6*-GFP. Values represent the mean ± SEM of at least 3 independent experiments. Significance was determined using one-way ANOVA with multiple comparisons (Brown-Forsythe and Welch’s corrected); **P < 0.01, ****P < 0.0001. (D) Quantification of sterol content in transformed ADΔΔ strains and BY4741 control. The ADΔΔ + *PDR6* displays increased filipin staining compared to the ADΔΔ control and ADΔΔ + *AFR1* but is indistinguishable from the WT BY4741 strain. Log-phase cells were incubated with filipin and imaged/quantified as described in Fig. 5. Values represent the mean ± SEM where n=50. Significance was determined using one-way ANOVA with multiple comparisons (Brown-Forsythe and Welch’s corrected); **P < 0.01.

### *PDR6* expression does not restore efflux activity but increases sterol staining in the ADΔΔ strain

Using the same set of strains as above, we measured FLC susceptibility with Etest strips. The strain expressing Afr1-GFP, a known FLC efflux pump, exhibited a 12-fold increase in MIC (MIC = 0.75 µg/mL) over the ADΔΔ+empty vector control (MIC = 0.064 µg/mL; Fig. 6B). In contrast, the ADΔΔ+Pdr6-GFP strain exhibited a modest increase with a MIC of 0.25 µg/mL, resulting in a 4-fold increase over the control but a 3-fold decrease compared to the ADΔΔ+Afr1-GFP strain. If this small increase in MIC was due to Pdr6-efflux activity, we should be able to measure it using the flow cytometry assay described above. As shown in Fig. 6C, no differences were seen in the amounts of accumulated drugs between the ADΔΔ+empty vector control and the strain expressing *PDR6*-GFP. In contrast, expression of *AFR1*-GFP resulted in a significant decrease in the levels of R6G and Probe 1, albeit not to the levels seen in the WT BY4741 control strain. The decrease in NR was not statistically significant, but a clear downward trend was also observed in the ADΔΔ+Afr1-GFP strain. Next, given the filipin-staining results shown in Fig. 5, we reasoned that the small MIC increase seen could be due to the potential role of Pdr6 in ergosterol transport. To test this, we subjected the set of ADΔΔ strains to filipin staining and analysis as above. Consistently, the ADΔΔ strains with an empty vector or expressing Afr1-GFP had the lowest PM fluorescence, while the ADΔΔ+Pdr6-GFP had fluorescence that was indistinguishable from the WT control (Fig. 6D). All of these results, together with the localization of Pdr6-GFP, suggest that the role of Pdr6 in azole resistance stems from effects on ergosterol synthesis and/or transport and not from acting as a specific FLC efflux pump.

### *PDR6* influences interaction with host macrophages

Alveolar macrophages represent the first line of defense against cryptococcal infections and the complex fungal-phagocyte interaction ultimately determines the course of infection and patient outcome (41). We initially identified *PDR6* in a screen for fungal regulators of phagocytosis (26). We used the same automated high-content imaging method to quantify the phagocytic index (PI) of the library’s *pdr6*Δ strain and the recreated strains by a human monocytic cell line (THP-1) (Fig. 7A and S1D). In agreement, the original and recreated *pdr6*Δ mutants were all phagocytosed more readily than the WT, providing evidence that changes in phagocytosis are a direct result of loss of *PDR6* function. The *pka1*Δ strain served as a high-uptake positive control (26). This observation begs the question of how the deletion of an atypical PDR-type ABC transporter was affecting the cryptococcal-host interaction. We first tested if the absence of *PDR6* affected the cryptococcal capsule. The *pdr6*Δ mutant was able to generate capsule comparable to the WT, indicating that *PDR6* is not involved in capsule formation (Fig. S2C). Next, we assessed if there were any defects in cell-wall structure by growing *pdr6Δ*, WT, *afr1*Δ, and *afr1*Δ/*afr2*Δ/*mdr1*Δ strains under cell-wall-stress conditions (Fig. S2D). All strains (except for *erg3*Δ) grew under the tested conditions, and melanized normally, suggesting that there are no defects in the cell wall (Fig. S2D,E). Finally, we hypothesized that *PDR6* was involved in the secretion/efflux of an immunomodulatory factor and thus influencing host interactions. To test this idea, we performed similar uptake assays as previously described, but used strain-specific conditioned media (CM) during the cryptococcal-macrophage coincubation. The thought process being that if *PDR6* is in fact responsible for the release of an immunomodulatory compound, then the CM from the *pdr6*Δ strain should influence the uptake of the WT as to increase the PI values, whereas the CM from the WT should decrease the PI values of the *pdr6*Δ strain. As shown in Fig. 7B, there were significant differences in phagocytosis dependent on the source of the CM. Specifically, the incubation of the WT with *pdr6*Δ CM increased uptake ∼2-fold, while the incubation of the *pdr6*Δ strain with WT CM decreased uptake back to WT levels. Notably, use of CM from the *afr1*Δ mutant had no significant impact on the PI values of either WT or *pdr6*Δ strains. To test if this effect was specific for cryptococcal cells or impacted all phagocytic cargo, we carried out the same assay with non-pathogenic *S. cerevisiae* (Fig. 7C). The host cells were able to phagocytize a significant portion of this yeast regardless of the CM used. This suggests that *PDR6* is involved in the release of an immunomodulatory factor that is present in the media and significantly impacts phagocytosis of cryptococcal cells by host phagocytes. Taken together, these data confirm that the *pdr6*Δ mutant is recognized and phagocytosed more avidly than the WT strain, and that the deletion of *PDR6* alters the secretion/efflux of cryptococcal immunomodulatory compounds.

**FIG 7.**
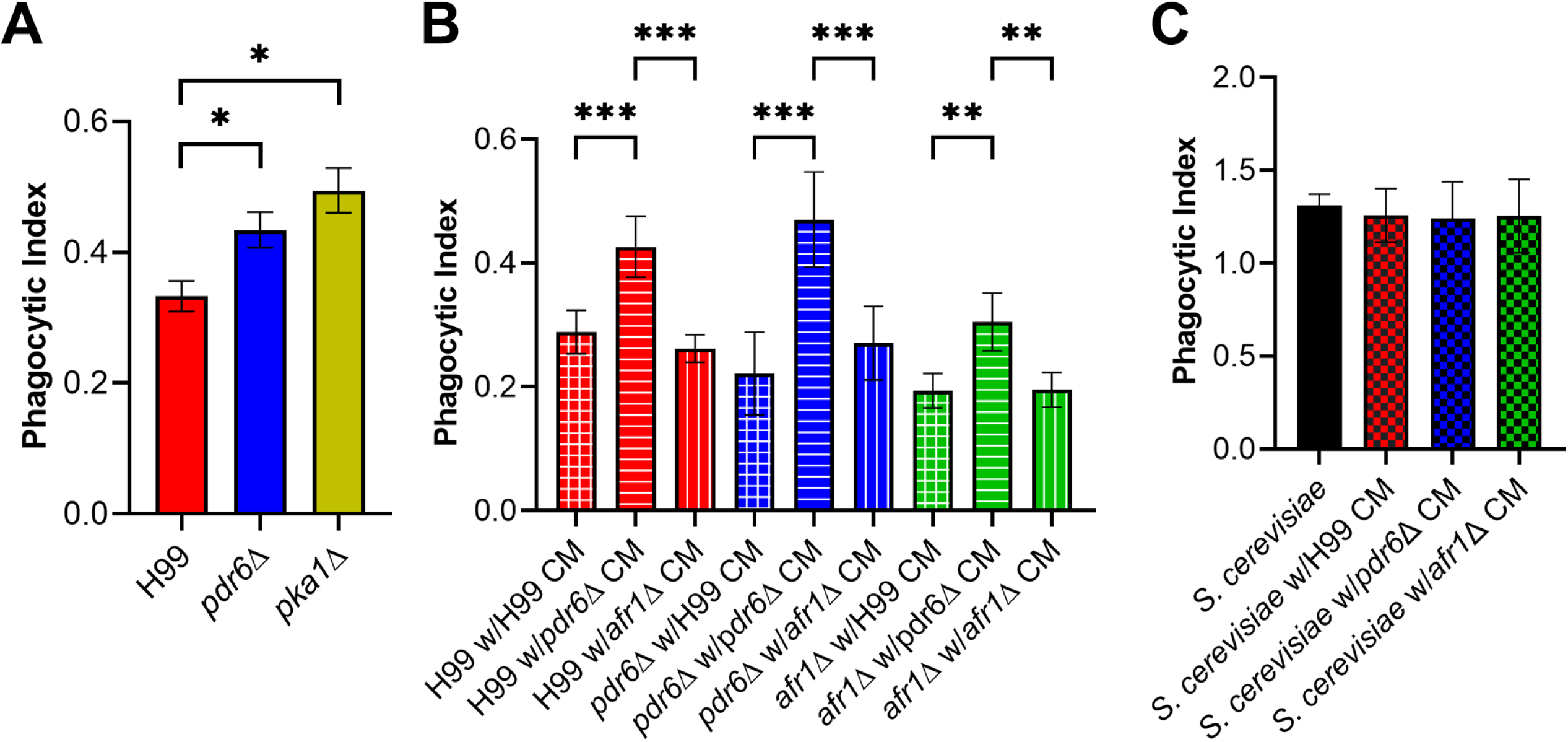
The *pdr6*Δ mutant conditioned media (CM) increases the phagocytosis of other cryptococcal strains by host macrophages. (A) Uptake assays of WT and deletion strains were performed to determine phagocytic indices of respective strains. The *pdr6*Δ mutant displayed increased uptake compared to the H99 strain. The *pka1*Δ cells were used as a positive control. Stained and opsonized log-phase cells were incubated for 1 hr with THP-1 cells and imaged using automated microscopy. Values represent the mean ± SEM from three independent experiments. (B) Uptake assay using CM from cryptococcal strains was performed. CM obtained by centrifugation and filtration of supernatants from 5-day cultures incubated at 37°C + 5% CO_2_ in RPMI-1640 media. The *pdr6*Δ CM altered the phagocytic uptake values of the WT strain and vice versa, whereas the *afr1*Δ CM has no effect. Values represent the mean ± SD of a representative experiment from three independent experiments. (C) Uptake assays with CM were performed as in (B) but using *S. cerevisiae* strain BY4741. BY4741 is easily phagocytized by THP-1 cells under all conditions. Significance was determined using one-way ANOVA with multiple comparisons (Brown-Forsythe and Welch’s corrected), *P<0.05; **P<0.01; ***P<0.001.

### *PDR6* is involved in capsular shedding and adherence

To investigate in more detail how *PDR6* is altering the cryptococcal-phagocyte interaction, we focused on the polysaccharide capsule. Primarily composed of glucuronoxylomannan (GXM) and glucuronoxylomannogalactan (GXMGal), the capsule is known to be secreted/shed in a constitutive manner. This secreted capsule plays a central role in virulence by acting as an antiphagocytic factor, providing protection against host stressors, and modulating the host immune response (42, 43). We wondered if the release/secretion of the capsule into the surrounding environment is altered in the *pdr6*Δ mutant. To test this, we analyzed the amount of shed capsule by electrophoresis and immunoblotting using anti-GXM antibodies (Fig. 8A) (44). Quantification of the immunoblots (Fig. 8A and S2G) showed that the *pdr6*Δ mutant sheds less capsule compared to the WT in both DMEM and YNB media, and this was significantly more pronounced in DMEM (Fig. 8A). Notably, there were no differences in shed capsule between the *afr1*Δ, *afr1*Δ/*afr2*Δ/*mdr1*Δ, or *erg3*Δ strains and the WT, indicating that this phenotype is unique to the Pdr6 transporter and not related to low levels of ergosterol. The acapsular strain *cap59*Δ was used as a control. This observed defect in capsule shedding could explain the phagocytosis phenotype, but to further test if there are functional consequences associated with this defect, we performed biofilm/adherence assays (Fig. 8B). Previous studies have shown that shed GXM is essential for the formation of cryptococcal biofilms (45–47). Consistently, we found a similar trend where the *pdr6*Δ strain formed weaker/less biofilm structures when compared to the WT, as measured by biomass quantification using anti-GXM antibody. Identical results were obtained by metabolic quantification using XTT reduction assays (data not shown). Taken together, these data suggest that *PDR6* is implicated in the shedding of the polysaccharide capsule, affecting both phagocytosis by host cells and biofilm formation.

**FIG 8.**
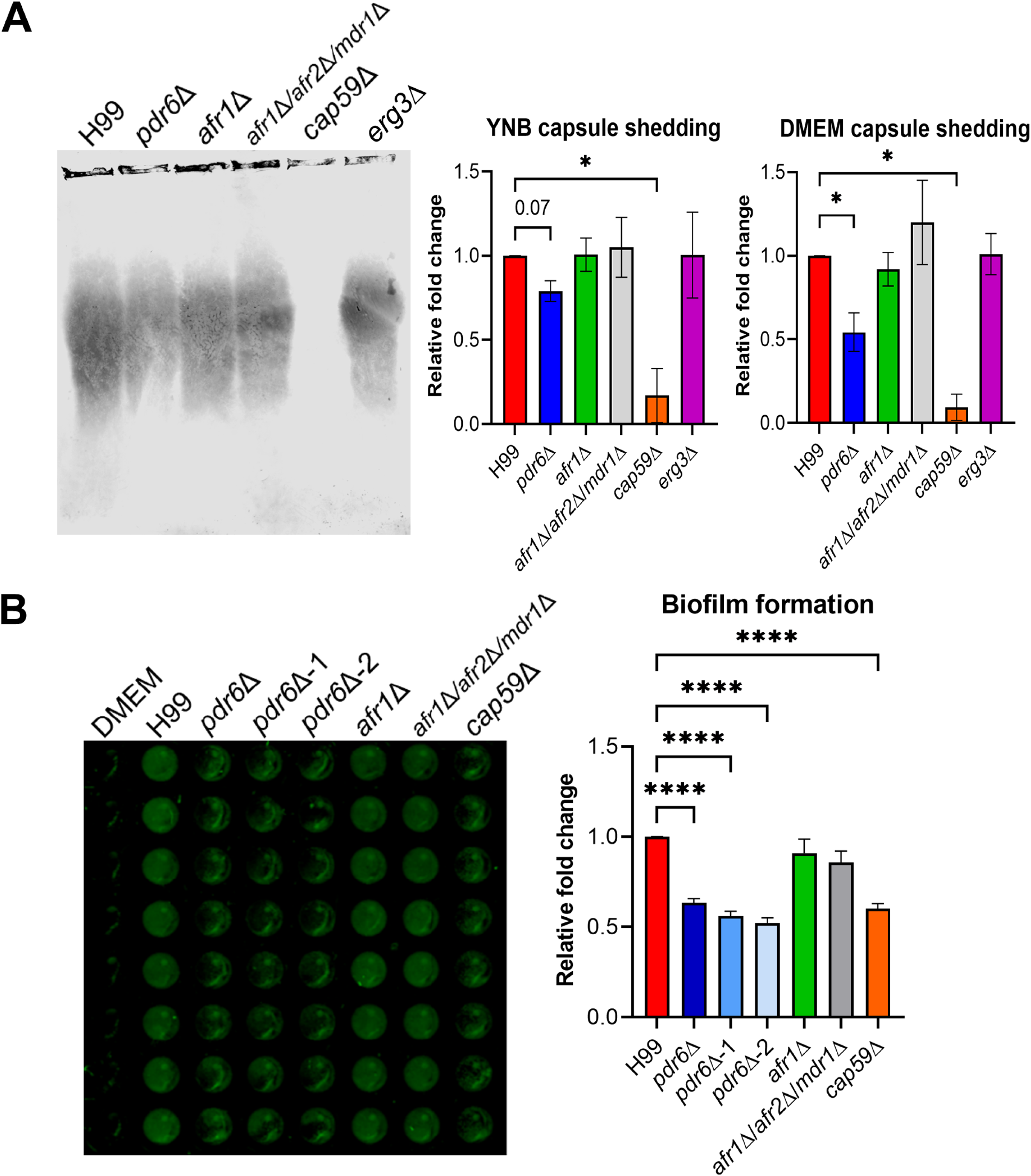
Deletion of *PDR6* alters capsular secretion and biofilm formation. (A) Capsule shedding assays were performed to determine amount of GXM secreted from cryptococcal strains into the media. (Left) Representative immunoblot showing shed GXM from cells grown for 2 days in YNB. Visualization of shed GXM was done using fluorescent detection (Odyssey, LiCor). Quantification of signal revealed that the *pdr6*Δ mutant shed significantly less capsule than WT under DMEM (host-like) conditions and, albeit not statistically significant, there was a clear trend of less shedding in YNB conditions as well. Values from at least 3 independent experiments were normalized to the WT shed capsule and combined. Bars represent the mean ± SEM. (B) Biofilm-forming assays were performed in nontreated microtiter plates, incubated for 72 hr, washed, and visualized for biofilms using a fluorescent imaging system (Odyssey, LiCor). Quantification of signal reveal that all *pdr6*Δ mutants form weaker/less adherent biofilms when compared to the WT. Values from at least 3 independent experiments were normalized to WT and combined. Bars represent mean ± SEM. Significance was determined using one-way ANOVA compared to H99 (Brown-Forsythe and Welch’s corrected); *P<0.05; ****P<0.0001.

### *PDR6* function is required for *in vitro* and *in vivo* virulence

Given the altered host interactions exhibited by the *pdr6*Δ mutant, its mild thermotolerance (Fig. S2F), and the decreased capsule shedding, we next assessed its ability to survive within the challenging and harsh intracellular environment. To test this, we performed *in vitro* survival assays and found that the *pdr6*Δ mutant struggled to survive within host macrophages when compared to the WT (Fig. 9A). After 4h in the presence of naïve human macrophages, ∼10% of intracellular WT cells died, whereas 25%-35% of the *pdr6*Δ cells are killed. To expand on this mild *in vitro* defect, we tested the virulence of the *pdr6*Δ mutant in a murine inhalation model of cryptococcosis, monitoring disease progression by weight loss. Infection with the WT strain resulted in all animals steadily losing weight by about two weeks, 50% of them dying in 17 days, and all succumbing to infection by day 19 (Fig. 9B). In contrast, mice infected with the original *pdr6*Δ and the recreated *pdr6*Δ-1 strains lived about two weeks longer than the WT-infected mice, with animals beginning to die at day 34, and all succumbing to infection by day 41, thus showing a significant increase in survival. Notably, CFUs recovered from these mice at time of death showed significantly less organ burden compared to WT-infected mice, and a clear defect in dissemination to brain and spleen (Fig. 9C), supporting an altered progression or attenuated pathology. Consistently, the *pdr6*Δ-infected mice exhibited respiratory symptoms (rapid and labored breathing) that were absent in the WT-infected mice, and their lungs were significantly more inflamed and enlarged than the WT lungs at time of death. Taken together, these data suggest that *PDR6* function is required for full cryptococcal pathogenesis, and in its absence, the manifestation of the disease is protracted with altered symptomatology.

**FIG 9.**
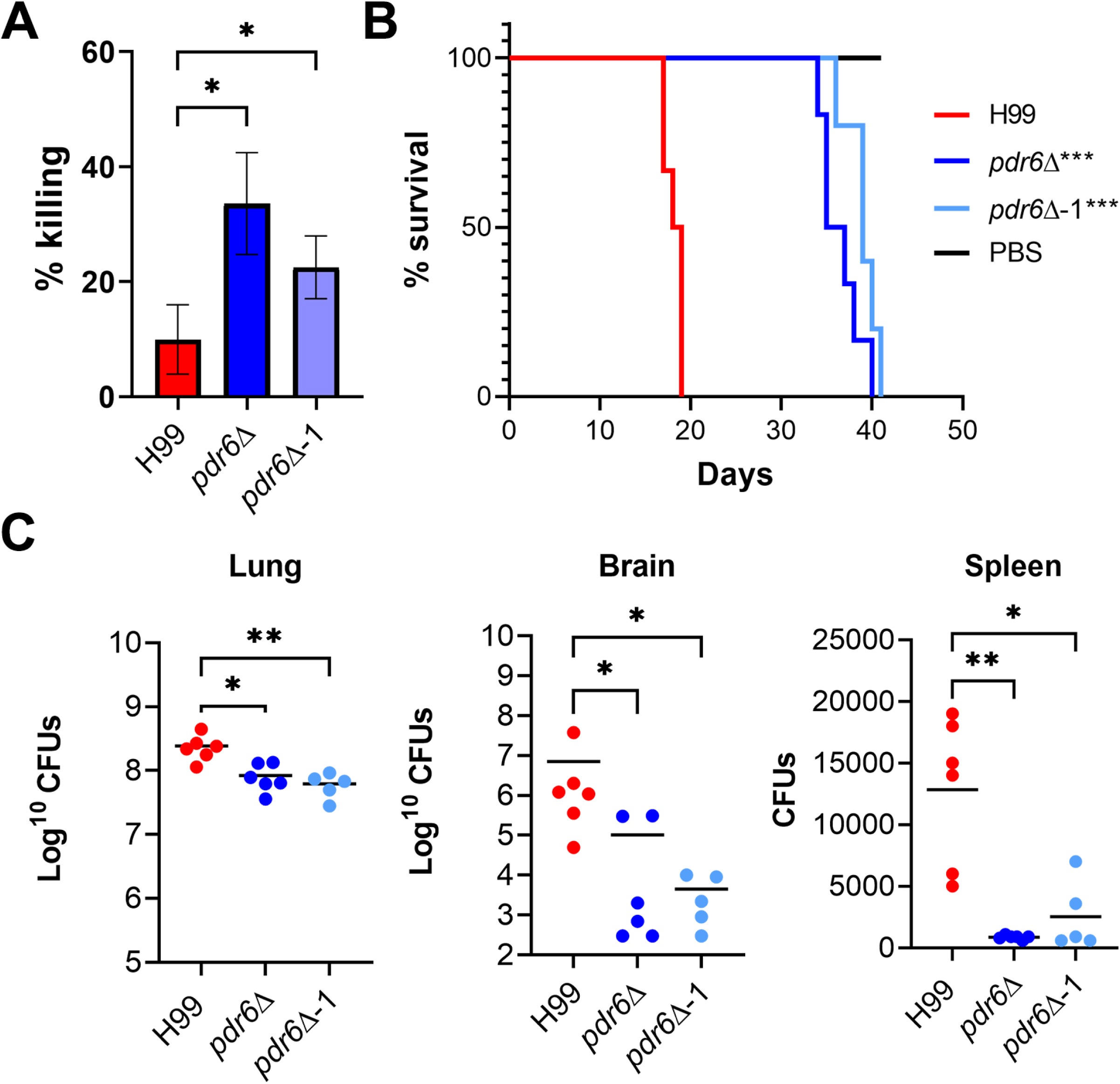
*pdr6*Δ mutant displays attenuated virulence *in vitro* and *in vivo.* (A) *In vitro* survival of H99 and *pdr6*Δ strains. Fungi and THP-1 cells were coincubated for 1 hr at an MOI of 0.1, washed to remove free cryptococci, and half the samples lysed to determine intracellular fungi and the other half grown for an additional 3 hr, after which they were also lysed. Shown in % killed, values represent the mean ± SEM from five independent experiments. Significance was determined using RM one-way ANOVA compared to H99 (Geisser-Greenhouse correction); *P<0.05. (B) *In vivo* survival curve of H99 and *pdr6*Δ-infected mice. Five to six AJ/Cr mice per group were infected intranasally with 5 x 10^4^ *C. neoformans*, or mock infected with DPBS and monitored for up to 41 days (experiment terminated when the last infected mouse died). Significance was determined using Mantel-Cox test; ***P<0.001. (C) Calculated organ burden of H99 and *pdr6*Δ strains at time of death. Organs were harvested, homogenized, and dilutions were plated on YPD. Plates were incubated for 48 hr and CFUs counted. Each data point represents a mouse and black bars represent the mean. There was significant less fungal burden in the lungs and a clear defect in dissemination. Significance was determined using a nonparametric one-way ANOVA with multiple comparisons (Dunn’s corrected), *P<0.05.

## DISCUSSION

The H99 genome contains 10 genes encoding PDR-type ABC transporters (Fig. 1B and Supplemental Text S1). Although it has a conserved representative gene in all clades except clade II, few have been characterized. Here we present an initial characterization of an atypical, half-size, PDR transporter found in a fungal-host interaction screen. This gene, which we call *PDR6*, is conserved in the fungal kingdom (clade VIb), but no function has been reported for that subgroup. Since all the other members of that clade are pathogenic fungi, our work may have broad implication for medically-important fungi. Given the similarity of *PDR6* to *AFR1* and *PDR5* (*AFR2*), which have been shown to act as efflux pumps for FLC, we tested the hypothesis that *PDR6* might act in a similar way. However, our data show that it is unlikely that Pdr6 is a major efflux pump for the compounds tested, including FLC. The *pdr6*Δ mutant accumulates the same level of all these compounds, yet it is hypersensitive to them.

We then focused on ergosterol, the direct or indirect target of most antifungal drugs. Studies in the model yeast *S. cerevisiae* have documented the ergosterol biosynthetic pathway in detail (48). This occurs in the ER, yet how ergosterol is trafficked to the PM is not known, although it is known that vesicular trafficking is not involved (49). In this yeast, *PDR18* has been implicated in that process, and indeed, when *PDR6* is compared to the *S. cerevisiae* genome, *PDR18* is the gene closest to it, although there is no real *PDR6* homolog in *S. cerevisiae*. Moreover, fungal and plant PDR genes are part of the subfamily G of ABC transporters. In mammals, all ABCG members are half-size, and the mouse ABCG3 and human ABCG5 and 8 transporters are all involved in cholesterol transport (50). Interestingly, a BLAST analysis of these ABCG sequences against the *Cryptococcus* genome identifies *PDR6* as the gene with the highest similarity to ABCG3 and 8. Hence, our finding that Pdr6-GFP localizes to the ER and to the PM, the biosynthesis site and final destination of ergosterol, respectively, and that the *pdr6*Δ mutant has decreased ergosterol while the ADΔΔ+Pdr6-GFP increases PM ergosterol, strongly support a model were Pdr6 functions as an ergosterol transporter. Notably, this function seems not to be conserved in other PDR genes, as the *afr1*Δ and *afr1*Δ/*afr2*Δ/*mdr1*Δ mutants have normal levels of ergosterol as quantified by filipin staining. Interestingly, the *pdr6*Δ mutant is not more resistant to AMB as would be expected for strains lacking ergosterol (30), at least as judged by MIC values. A previous study of the susceptibility of young and old cells to AMB found that cells lacking *PDR6* are less resilient to AMB treatment, but this was measured as % survival and not as MIC values, preventing comparison of those results with ours (51). Overall, our data indicate that the diminished levels of ergosterol are low enough to promote sensitivity to azoles, but high enough to still be targeted by AMB and exert its antifungal activity.

It is known that defects in the PM can affect the cell wall and/or capsule (52, 53). We did not find evidence for gross alterations to either of these layers, but the defect in capsule shedding, and the results of the CM studies, are highly suggestive of Pdr6 having a role in regulated shedding of the capsule. It has been shown that capsule is shed constitutively, but recent studies have demonstrated that capsule shedding can also be regulated (43). *PDR6* is highly and rapidly induced under host conditions, which are conditions known to induce capsule growth and shedding. Notably, it is known that normal levels of ergosterol are needed for thermotolerance, hence if *PDR6* is involved in ergosterol homeostasis, it would be expected to be induced upon shift to body temperature. The altered ergosterol levels in the *pdr6*Δ mutant could explain both the increased sensitivity to FLC and other azoles, and the mild defect in thermotolerance (Fig. 10). However, low ergosterol cannot explain the reduced capsule shedding in this mutant as the *erg3*Δ mutant, which also has low levels of ergosterol, sheds similar amounts as WT. Additionally, other ergosterol mutants have also been shown to shed normal levels of capsule (30). This phenotype still needs more investigation.

**FIG 10.**
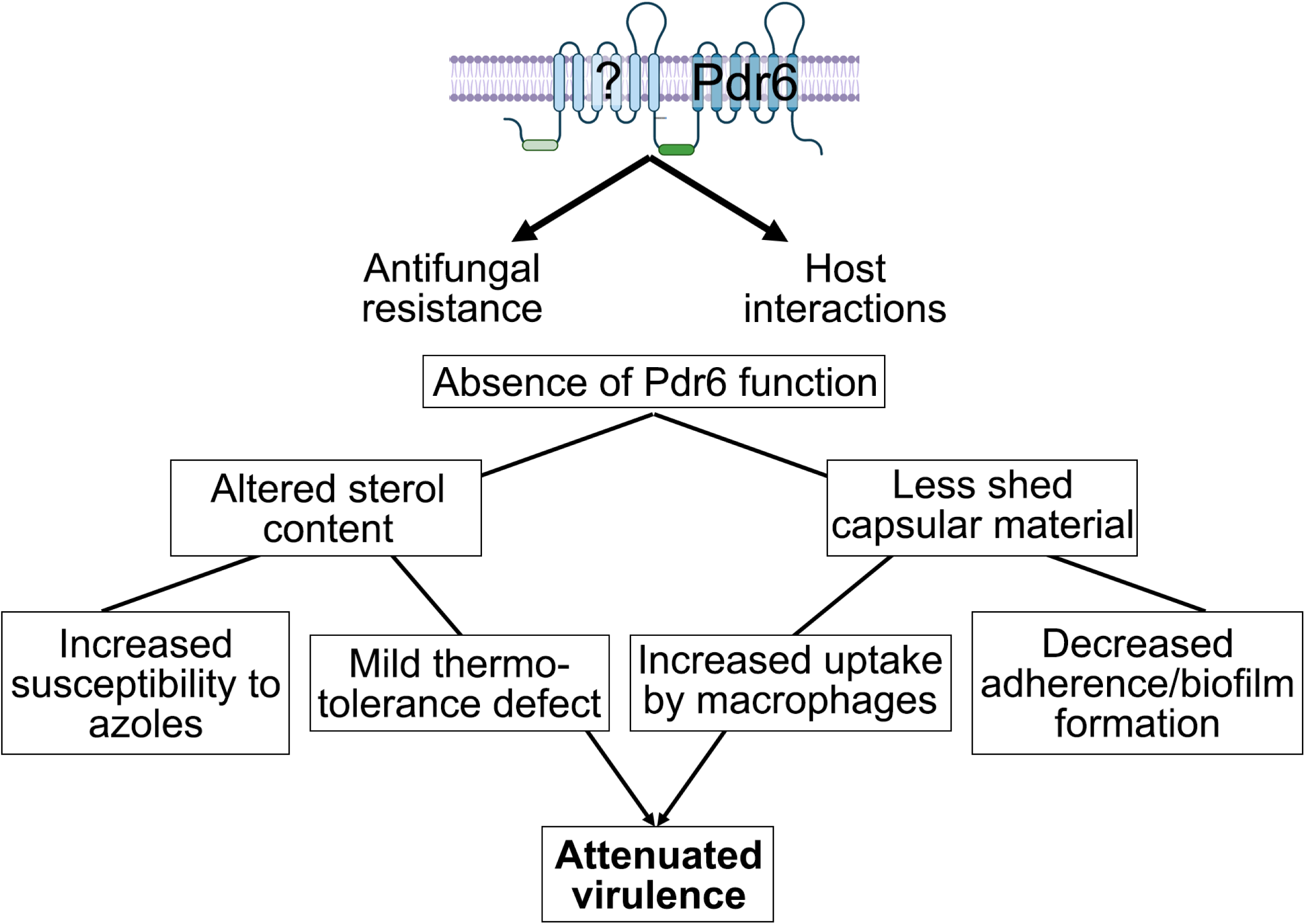
Model of Pdr6 and relationship to antifungal resistance, host interactions, and virulence. Pdr6 is a half-size transporter (dark blue), hence it needs to dimerize with itself or another half-size transporter (light blue) to be functional. In its absence, altered sterol content contributes to azole and thermal susceptibility, and the reduced capsule shedding promotes recognition and phagocytosis by host macrophages. Together, these phenotypes result in altered pathogenesis and attenuated virulence.

Lastly, we show that Pdr6 function is needed for normal progression and disease manifestation in the mouse inhalational model. This could be explained by the decreased capsular shedding, high phagocytosis, and lower intracellular survival (Fig. 10). The increased recognition and phagocytosis could give the host an advantage since the fungus does not survive as well intracellularly. This would affect dissemination, as there is ample evidence that phagocytes are needed for extrapulmonary dissemination, a process known as trojan-horse transit (54). However, in order for trojan-horse transit to occur, the fungus must be able to survive intracellularly, and the *pdr6*Δ mutant is defective in this respect. Moreover, the decreased capsule shedding from the extracellular fungi would result in a stronger inflammatory response. The *pdr6*Δ mutant is also capable of inducing production of a large capsule, hence over time, less cells would be phagocytosed, and free cells in the tissue would accumulate. With the increased fungal burden, and the stronger inflammatory response, the host advantage is lost, and the animals die exhibiting different symptoms. Ongoing studies into the immunological response of the animals to *pdr6*Δ infection will shed more light on this hypothesis.

In conclusion, we have described a new mechanism by which cryptococcal cells can modulate their susceptibility to azoles: ergosterol transport regulated by Pdr6. Our results with the CM also indicate a potential new antiphagocytic mechanism: the *pdr6*Δ mutant can still grow a capsule, yet it is recognized and phagocytosed more avidly. Given that there are *PDR6* homologs in other fungal pathogens, it would be interesting to see if this function is conserved. Notably, a *PDR6* ortholog is missing in *C. deneoformans*, but present in *C. gattii*. *C. deneoformans* is considerably less pathogenic than *C. neoformans,* and *C. gattii* is considered a primary pathogen, although it is more associated with pneumonia than CNS infection. Would absence of *PDR6* in *C. deneoformans* contribute to the virulence differences? Moreover, in addition to *PDR6*, *C. neoformans* has two additional half-size PDR transporters, which are missing in *C. gattii*. These are not in clade VIb, but their function is nevertheless unknown and could be related to Pdr6 since, in order to form a functional transporter, Pdr6 must homo- or heterodimerize. The binding partner of Pdr6 might influence the function of the resulting complex, which could be important under certain environmental conditions, such as inside a host. Elucidation of the diverse functions of ABC transporters, specifically the PDR-type, has the potential to open new lines of investigation for better treatments for this devastating disease.

## MATERIALS & METHODS

### Strains, cell lines, growth conditions, and reagents

All *C. neoformans* strains used were in the serotype A strain H99α background. The original *pdr6*Δ strain was obtained from the Madhani deletion collection (55) and the recreated strains were made in this background by biolistics (see below for details). H99α *afr1*Δ and H99α *afr1*Δ/*afr2*Δ/*mdr1*Δ were a generous gift from Yun Chang (National Institutes of Health). The *S. cerevisiae* strain ADΔΔ and the plasmids pABC3 and pABC3-KLmGH were graciously shared by Richard Cannon (Otago University, New Zealand). Maintenance and growth conditions, as well as media details can be found in Supplemental Text S1.

The human monocyte cell line THP-1 (ATCC TIB-202) was grown in THP-1 complete media and differentiated with phorbol 12-myristate 12-acetate (PMA, Sigma) as described in (55). THP-1 cells were split every 3-4 days and new vials were thawed every month. Details on complete media preparation and differentiation into macrophages can be found in Supplemental Text S1.

All antifungals, xenobiotics, and other chemicals used were obtained through VWR or from Sigma. Details on the preparation of stock solutions, including solvent concentrations, as well as their storage information can be found in Supplemental Text S1.

### Fungal genome manipulation

Since complementation strains of the original *pdr6*Δ (from the Madhani collection) proved challenging to create, we used the split marker method (56) to delete the whole *PDR6* coding region in H99α after amplifying NAT resistance split marker fragments from genomic DNA of strain 1F1 (*pdr6*Δ) of the Madhani deletion collection. Three independent *pdr6*Δ mutant strains were generated to verify that all observed phenotypes are a direct result of the loss of *PDR6* (Fig. S2). Additionally, we performed whole genome sequencing on the generated *pdr6*Δ mutants to confirm proper genetic manipulation (data not shown).

### Drug susceptibility assays

The CLSI M27-A3 reference method was used to determine the MIC of each drug (57). Briefly, the compounds to be tested were prepared at 2X the final concentration from stock solutions in RPMI 1640 medium. These were dispensed in a 96-well plate with 2-fold serial dilutions. 500 cells in the same volume as the drugs were then added to each well. Plates were incubated at 37°C + 5% CO_2_ for 48 – 72 hr. Minimal inhibitory concentrations (MICs) were quantified by measuring the OD_600_ using a microtiter plate reader. For details on the compounds and concentration ranges tested see Supplemental Text S1.

The MIC of FLC was also determined on RPMI plates with Epsilometer test strips (Etest strips; Liofilchem Thermo). Approximately 5 × 10^4^ cells were plated on RPMI agar plates before application of the Etest strips. Plates were incubated at 30°C for 48 hr.

### Checkerboard assays

Checkerboard assays to assess the fractional inhibitory concentrations (FICs) for combinations of antifungal drugs was performed as previously described (58). Log-phase cultures were used and mixed with the antifungal stocks in RPMI 1640. For more details on the procedure, final antifungal concentrations, and calculations of the FIC index, see Supplemental Text S1. A FIC index value of ≤ 0.5 is considered synergistic, additive if the value is between 1.0 and 0.5, and antagonistic if the FIC index is ≥ 2.0.

### Flow cytometry analysis of the efflux of R6G, Nile Red, and Probe 1

Accumulation assays was performed as previously described (24, 34). Briefly, strains were grown overnight in YPD at 30°C with shaking. Log-phase cultures were incubated with the fluorescent reporters in PBS for 30 min at 30°C with shaking. The reactions were stopped by cooling on ice, and the mixture was diluted 40-fold in cold 1X PBS. The accumulation was immediately assessed by flow cytometry. For details on the concentration of the reporters, the lasers and filters used, and how the data was analyzed see Supplemental Text S1. Synthesis of Probe 1 was done in the Mobashery lab following published methods (34). For a detailed synthetic scheme and characterization of the product see Supplemental Text S1.

### Liquid Chromatography/Mass Spectrometry (LC/MS)

Log-phase cultures were incubated with FLC as above. A total of 200 μL was removed for CFU quantification and the rest was processed to make the lysate. The lysate was mixed with same volume of cold MeOH, vortexed, and spun down. The supernatant was mixed with the same volume of CH_2_Cl_2_, vortexed, and spun down. The organic (bottom) phase was transferred into a glass tube, the solvent was evaporated and the residues was resuspended in MeOH. At this point the samples were further cleaned by passing through a solid-phase extraction (SPE) column. The elutes from SPE were concentrated by speedvac, reconstituted in 50:50 water:MeOH, and analyzed using LC/MS. For details on the instrumentation, columns, buffers, flow rates, software, and mass-spec parameters see Supplemental Text S1.

### Phenotyping

Strains tested were grown overnight in YPD, diluted to an OD_600_ of 0.2, and grown for two doublings. The cultures were diluted to 1 x 10^7^ cells/mL and serially diluted (10-fold) and spotted (5 μL) onto YPD with 1 M NaCl, 0.005% SDS, 0.1% Congo Red, and 0.5 μg/mL caffeine. Plates were incubated at 30°C for 48 – 72 hr.

### Uptake Assay

Uptake assays were performed as previously described (26). Briefly, differentiated THP-1 cells were incubated with Lucifer Yellow-stained fungal cells that had been opsonized with 40% fresh human serum obtained from healthy donors (approved by the UND IRB committee as a non-human subject research procedure). Following a 1 hr incubation, the plates were washed using a microplate washer (405LS, Biotek, Winooski, VT), fixed with 4% formaldehyde, and stained with DAPI (Sigma) and CellMask Deep Red (Invitrogen). NaN_3_ in PBS was added to the plates and images were obtained using a Zeiss Axio Observer microscope, equipped with an automated stage. Each well was automatically imaged in a 3×3 grid and the images were analyzed using a Cell Profiler pipeline to determine phagocytic index (PI) values. For more details on these procedures, including the Cell Profiler pipeline, see Supplemental Text S1.

### Conditioned media

Conditioned media (CM) was generated as previously described (59). Briefly, overnight cultures were diluted to 1 × 10^5^ cells/mL in RPMI 1640 media and incubated for 5 days at 30°C. Cells were removed by centrifugation and the supernatant was filtered through a 0.25-μm filter.

### Capsule induction

Cells were grown overnight in YPD, washed with DMEM, and 1 × 10^6^ cells/mL were added to 24-well tissue culture plates. Plates were incubated at 37°C + 5% CO_2_ for 24 hr. Cell suspension was collected and washed, India ink was added, and samples were visualized on a Zeiss microscope. Images were analyzed with ImageJ (NIH) for capsule thickness.

### Capsule shedding assay

Capsule shedding assays were performed as previously described (43, 44). Briefly, strains were grown overnight in YPD at 30°C, washed, and diluted to 1 × 10^6^ cells/mL in DMEM. Following a 15 hr incubation at 37°C + 5% CO_2_, the samples were heated to denature enzymes, centrifuged to separate cells and supernatant, and then stored at 4°C. The samples were loaded on a 0.6% certified megabase agarose (Bio-Rad) gel and subjected to electrophoresis for 15 hr at 25 V. Gel contents were transferred to a positively charged membrane using a standard Southern blot protocol with 10X SSC. Following overnight transfer, the membrane was blocked for 48 hr in 1X TBS-5% milk and incubated for 1 hr in 1X TBST-1% milk with 1 µg/mL of anti-GXM monoclonal antibody. Membrane was rinsed three times in 1X TBST and incubated for 1 hr in 1X TBST-1% milk with Odyssey antibody at 1:10,000. Membrane was rinsed again three times in 1X TBST and imaged on the Odyssey (Licor).

### Biofilm/Adherence assay

Cryptococcal adherence assays were performed in nontreated 96-well plates (CLS3631, Corning). Briefly, strains were grown overnight in YPD, diluted to 1 × 10^7^ cells/mL in DMEM, and 100 μl of cell suspension was added to the wells. Plates were incubated at 37°C + 5% CO_2_ for 72 hr, washed with PBS, fixed with 4% formaldehyde, blocked with 1.5% BSA and 0.1% NaN_3_ in PBS, and incubated with anti-GXM primary and secondary antibodies. Plates were imaged on the Odyssey and quantified using ImageStudio software.

Biofilms were also quantified by the XTT reduction assay, as previously described (60, 61). For details on this assay see Supplemental Text S1.

### Filipin staining

Staining of membrane sterols was performed as previously described (36, 62). Briefly, log-phase cultures grown in YPD were stained for 5 min with 5 μg/mL of filipin (Sigma) and washed with PBS. After washing cells were immediately visualized with an inverted Zeiss microscope. Fluorescence was calculated using Cell Profiler software. Additionally, fluorescent levels were quantified with a fluorescent plate reader (Bio-Tek). Fluorescence was measured at the range of 360 to 470 nm wavelength and normalized by OD_600_.

### Mouse virulence studies

To test virulence in a murine model, strains were cultured overnight in YPD, collected, washed, and diluted to 10^6^ cells/mL in DPBS. 50 μL of the cell suspension (or DPBS for the mock-infected) was used to intranasally inoculate groups of five mice (5 to 6-week-old female A/Jcr mice; Jackson Laboratories). Animals were closely monitored and sacrificed if they lost >20% relative to peak weight or at the end of the experiment (41 days). Homogenates of lungs, brains, and spleens were plated to determine organ burden.

## ACKNOWLEDGMENTS

We acknowledge assistance and support for the LC-MS/MS by Dr. Mijoon Lee from the Notre Dame Mass Spectrometry Core. We thank Dr. Richard Cannon for their gift of the ADΔΔ strain and associated plasmids. We acknowledge financial support from the Warren Center for Drug Discovery for a Sean Cocchia Rare Disease Research seed grant. We also thank members of the Santiago-Tirado lab for thoughtful comments and feedback on this work.

## FIGURE LEGENDS

**TEXT S1** Supplemental methods and details on naming nomenclature for PDR genes in *C. neoformans*.

**FIG S1** Antifungal and phagocytosis phenotypes in original and recreated *pdr6*Δ mutant strains. (A) Plates with FLC Etest strips to assess the zones of inhibition of H99 WT and *pdr6*Δ mutant cells grown on standard (pH 5.5) or alkaline (pH 7.5) YNB medium. Regardless of pH, the *pdr6*Δ mutant was 4X more susceptible than WT. The H99 and *pdr6*Δ mutant displayed FLC MICs of 64 and 16 μg/mL on the 5.5 pH plates and 24 and 6 μg/mL under alkaline conditions. Plates were imaged after 72 hr of growth at 30°C. (B) MIC of FLC for H99 WT and *pdr6*Δ strains were determined by broth microdilution assays after 48 h of growth at 30°C in standard and alkaline YNB. Microdilution assays were repeated three times for each condition and black bars represent the average MIC respectively. (C) Broth microdilution assays were used to compare FLC MICs of three independently recreated *pdr6*Δ strains to the *pdr6*Δ strain from the deletion collection. Assay was repeated three times and black bars represent the average MIC. (D) Comparative phagocytic indices of the library and recreated *pdr6*Δ mutants with the H99 strain. All *pdr6*Δ strains displayed significantly higher phagocytic values than the WT. Values represent the mean ± SD of a representative experiment from three biological replicates. Statistical significance determined using one-way ANOVA with multiple comparisons (Brown-Forsythe and Welch’s corrected), *P<0.05, **P<0.01.

**FIG S2** *PDR6* expression and virulence traits. (A) Transcript levels of *PDR6* in response to FLC exposure as measured by qRT-PCR. Log phase cells were either untreated (control) or treated with 4 μg/mL of FLC for 2 hr. No significant difference was observed in the presence of the antifungal. Expression levels were normalized to that of the actin gene. Values represent the mean ± SEM of three biological replicates. (B) Data from an RNA-seq experiment to analyze expression levels of *PDR6* after shifting to host-like conditions. Samples were taken prior to incubation (time = 0), and at 90, 180, 480, and 1,440 minutes after. Expression increased almost 3-fold in the first 90 min and remained relatively high throughout 24 hr. (C) Capsule induction assays were performed to determine capsule thickness in cryptococcal strains. Visualization and quantification of 24 hr cultures revealed that there was no significant difference in capsule size between the H99 and *pdr6*Δ mutant. Scale bar, 5 μm. Values represent the mean ± SEM where n=50. (D) Growth on stress plates was assessed to determine potential membrane or cell wall defects in the indicated cryptococcal strains. No differences were observed in the *pdr6*Δ mutant. Mid-log phase cultures were serially diluted on plates with cell wall/PM stressors and incubated for 72 hr at 30°C. CR, Congo Red. (E) Melanization was assessed by appearance of pigmentation on caffeic acid plates. 5.0 x 10^3^ cells were spotted and incubated as above. (F) Thermotolerance was assessed by spotting strains on YPD plates and incubating plates at 30 °C and 37 °C for 72 hr. A mild defect in growth was observed in all the *pdr6*Δ mutants except *pdr6*Δ-3. (G) Representative immunoblot showing shed GXM from cells grown for 24 hr in DMEM, 37 °C and 5% CO_2_.

## Notes

### Competing Interest Statement

The authors have declared no competing interest.

