## Supplementary figures and images for "An atypical ABC transporter is involved in antifungal resistance and host interactions in the pathogenic fungus *Cryptococcus neoformans*"

### Supplemental Fig. S1

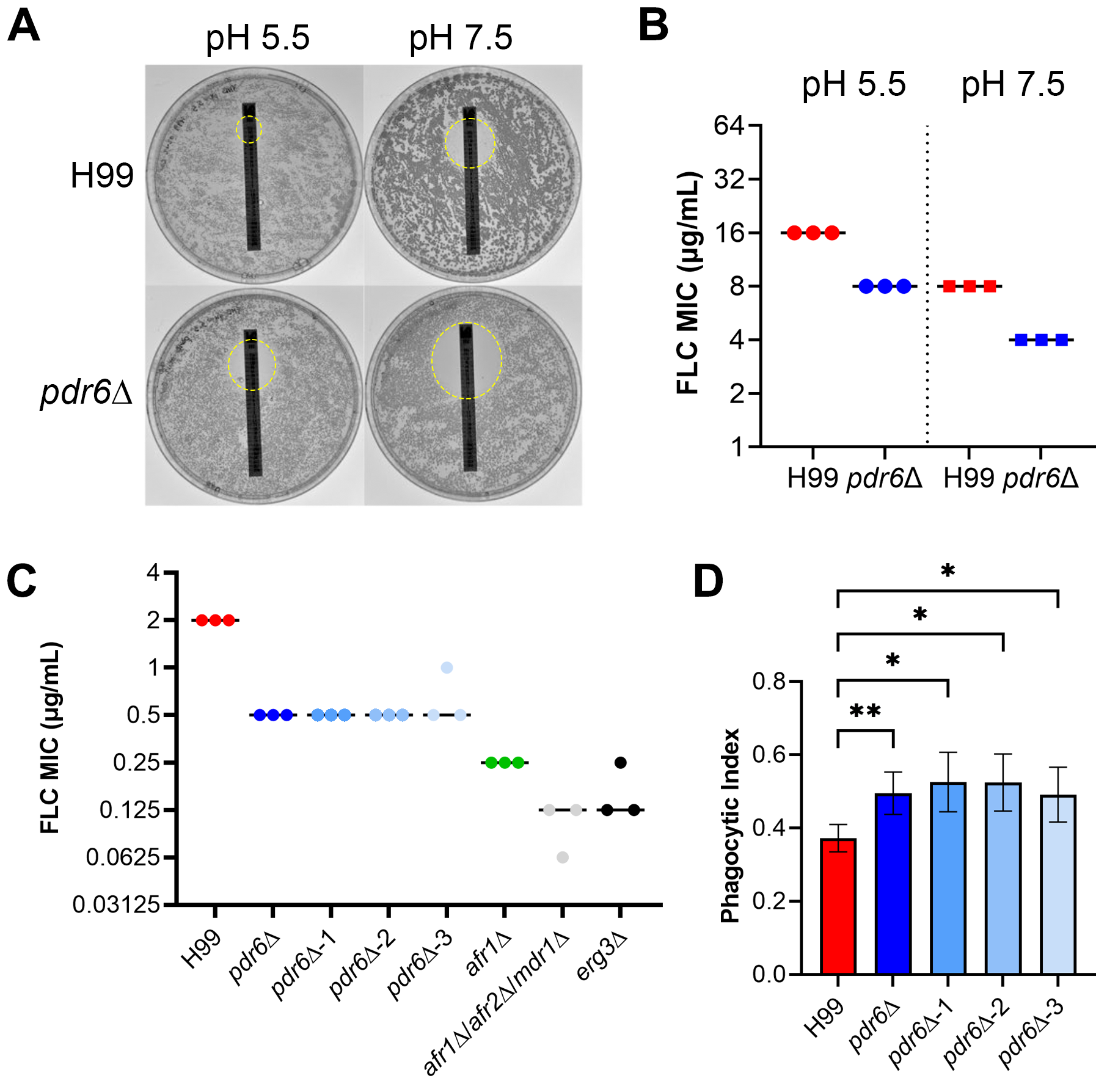

### Supplemental Fig. S2

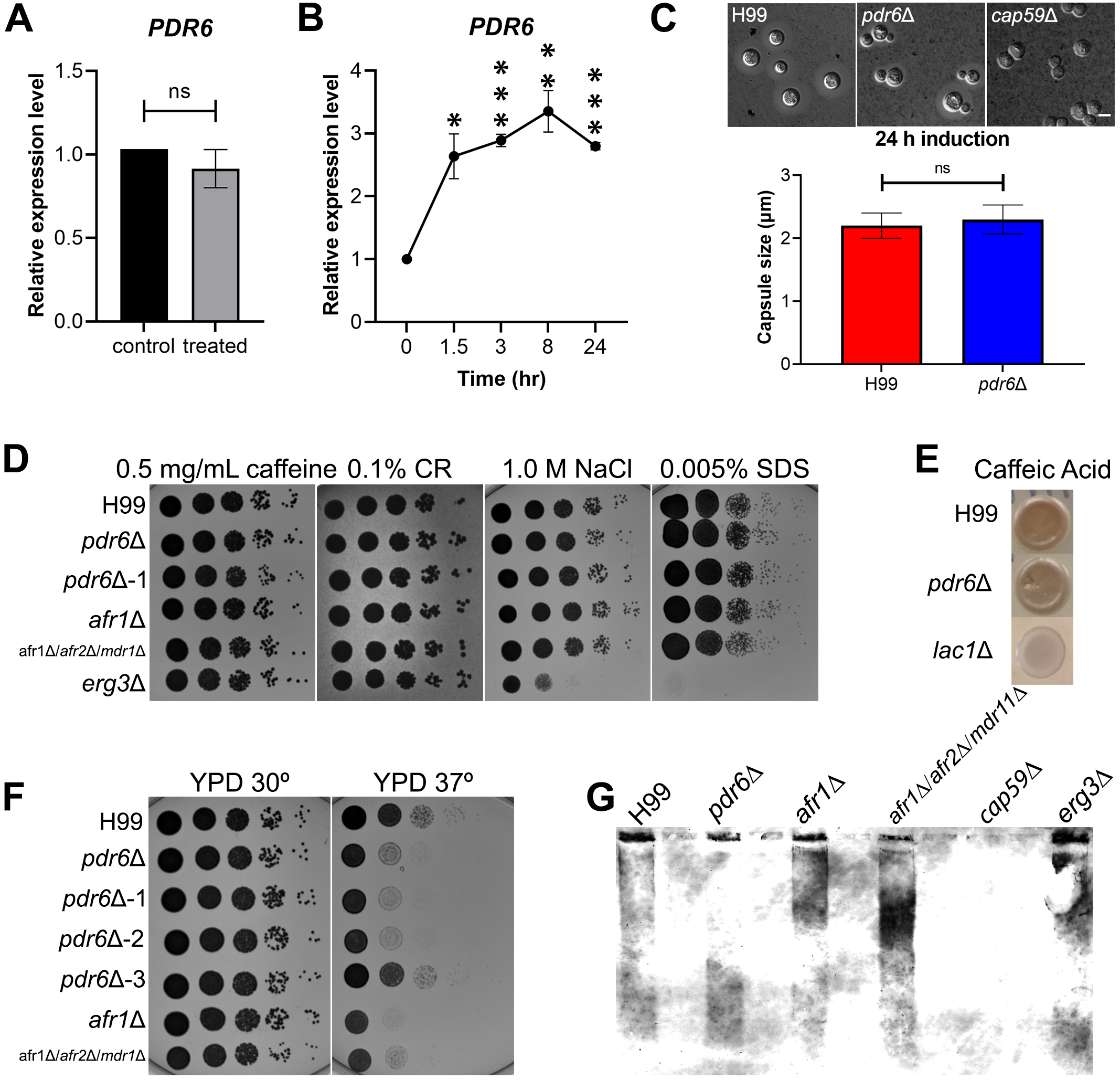
